# Efficient image analysis for large-scale next generation histopathology using pAPRica

**DOI:** 10.1101/2023.01.27.525687

**Authors:** Jules Scholler, Joel Jonsson, Tomás Jordá-Siquier, Ivana Gantar, Laura Batti, Bevan L. Cheeseman, Stéphane Pagès, Ivo F. Sbalzarini, Christophe M. Lamy

**Affiliations:** Wyss Center for Bio and NeuroEngineering, Geneva, Switzerland; Technische Universität Dresden, Faculty of Computer Science, 01069 Dresden, Germany; Max Planck Institute of Molecular Cell Biology and Genetics, 01307 Dresden, Germany; Center for Systems Biology Dresden, 01307 Dresden, Germany; Center for Scalable Data Analytics and Artificial Intelligence ScaDS.AI, Dresden/Leipzig, Germany; Unit of Anatomy, Faculty of Medicine, University of Geneva

## Abstract

The large size of imaging datasets generated by next-generation histology methods limits the adoption of those approaches in research and the clinic. We propose pAPRica (pipelines for Adaptive Particle Representation image compositing and analysis), a framework based on the Adaptive Particle Representation (APR) to enable efficient analysis of large microscopy datasets, scalable up to petascale on a regular workstation. pAPRica includes stitching, merging, segmentation, registration, and mapping to an atlas as well as visualization of the large 3D image data, achieving 100+ fold speedup in computation and commensurate data-size reduction.

## Main

The microscopic analysis of morphological and molecular markers in healthy and pathological organs is an essential component of medical diagnosis and biomedical research. This approach is usually carried out in small tissue sections of a few micrometer thickness, thus introducing a sampling bias in the assessment of biological and pathological processes. Next generation histopathology methods propose to analyze large organ parts at high resolution thanks to the use of new tissue preparation protocols and large-scale volumetric microscopy techniques^1–4^. Their wider adoption in research and in the clinic, however, is currently limited by the lack of appropriate methods to efficiently process and analyze the terabyte-sized datasets they generate. This means that currently several weeks of computer time are required between imaging and analysis, even on costly parallel distributed-memory computer clusters or graphics processing units (GPUs), prohibiting repetition of experiments on statistically significant cohorts or the achievement of acceptable turnaround times in clinical pathology.

To address this issue, we developed a complete image analysis pipeline based on the Adaptive Particle Representation (APR)^5^, which enables efficient analysis of large histology datasets, scalable up to petabytes on a regular workstation (see methods). The proposed pipeline allows real-time processing for fundamental tasks such as stitching and cell segmentation, and it permits researchers to visualize and gain insight into their data already during the acquisition.

The APR works by locally adapting the voxel size to the information content, effectively downsampling flat regions (e.g. background or uniform foreground) which carry little information. In fluorescence microscopy, such regions typically cover the vast majority of voxels. Thus, by representing them at lower resolution, the APR enables data size reduction up to several orders of magnitude^5^. By representing the image using much fewer particles than original voxels, APR-native downstream image processing leads to significantly accelerated computing times and reduced hardware requirements. This has been shown for e.g. graph-cut segmentation^5^ and spatial convolutions^6^. However, key APR-native processing algorithms and visualization methods, especially for large multi-tile acquisitions, have been unavailable until now.

Here, we present a complete suite of algorithms tailored for the analysis of large, multi-tile next-generation histology images. It includes a conversion module, allowing each tile to be converted to its APR representation immediately upon acquisition, with conversion parameters determined automatically (see methods). Subsequent processing steps either directly operate on the APR representation, or on reconstructed local voxel patches, resulting in accelerated processing and vastly reduced memory footprints. This includes stitching, segmentation, merging, atlas registration, and mapping.

To demonstrate the possibilities offered by the proposed pipeline, we applied it to a full mouse brain dataset (2 channels, 9×8 tiles) acquired on a clarity-optimized light sheet microscope (COLM)^7^ (Fig. 1, Supp. Table 1). Compared to voxels, the APR representation of the image required 115 times less storage (Supp. Table 1). The signal channel, corresponding to genetically-tagged parvalbumin positive (PV+) neurons, was stitched using the proposed pipeline in 55 seconds and reconstructed by taking the maximum values on overlapping areas (Fig. 1e-g, Supp. Table 1). The pipeline was able to correct for motor position inaccuracies and optical misalignment, yielding comparable results to TeraStitcher^14^ in a fraction of the time (>200 times faster in the presented sample, see Extended Data Fig. 1).

**Figure 1:**
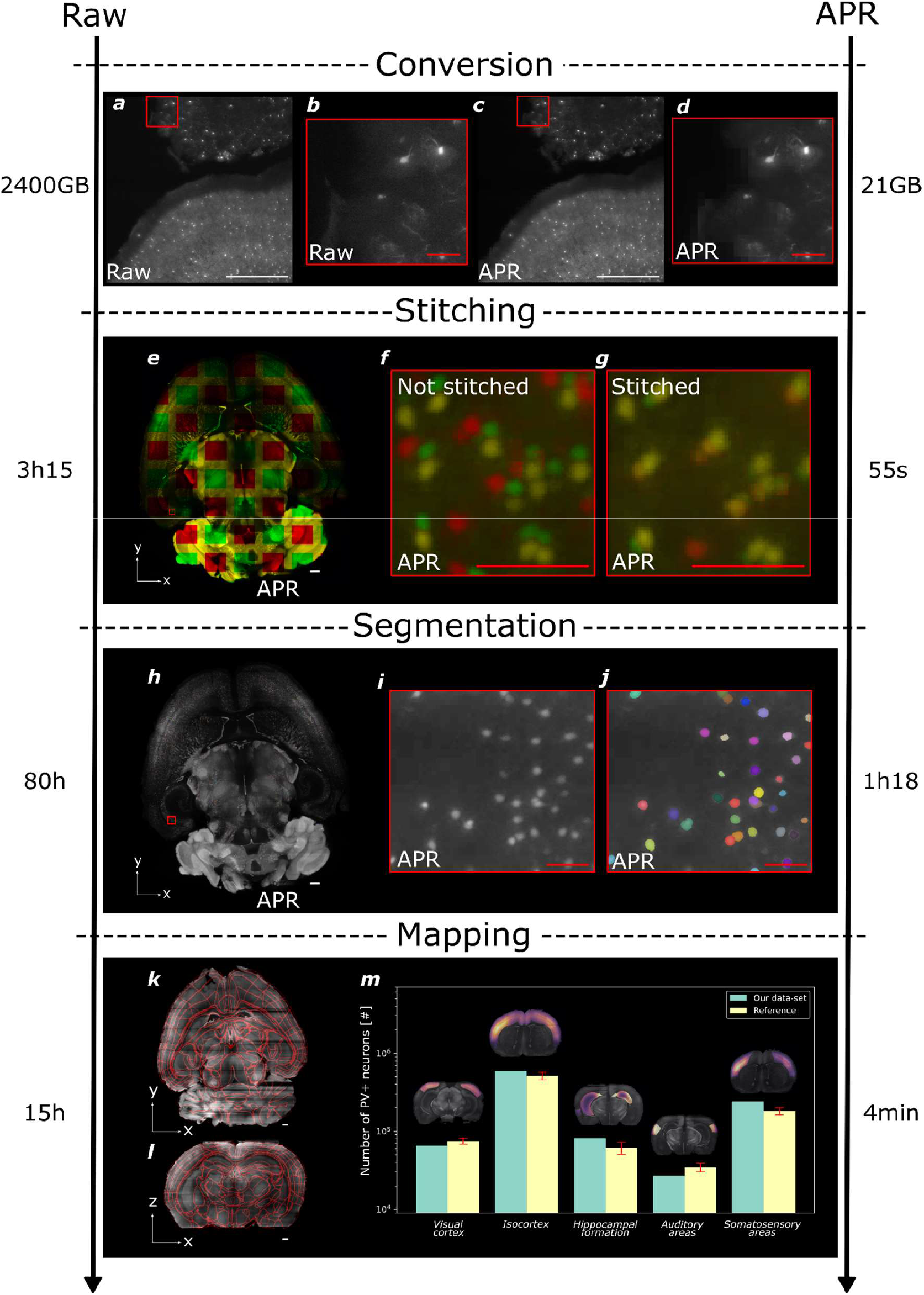
APR-native image analysis pipeline. **a**, Larger field of view of (b). **b**, Microscopic image of PV+ neurons in a cleared mouse brain, Raw data. **c,** APR converted data of (a) for a single tile. **d**, Magnified view of (c) matching (b). **e**, Horizontal view of the brain after stitching. Tiles are displayed in green and red. Overlapping areas appear in yellow. **f**, Magnified view of (e) before (using microscope position) and **g**, after stitching, showing alignment artifacts. The image shows a region where 4 tiles overlap. **h**, Horizontal view of the brain with segmented (random colored) PV+ neurons. Red square indicates subset displayed in (i) and (j). **i,** Magnified view of (h) (maximum intensity projection of 20 planes) before and **j**, after segmentation. **k,** Horizontal and **l**, coronal views of the brain autofluorescence channel overlaid with the Allen Brain Atlas region boundaries after registration. **m**, Comparison between the number of segmented PV+ neurons found in our dataset with a previously published reference^10^. For each brain region, the PV+ neuron density is overlaid as a color map (from purple for low density to bright orange for high density) on the acquired data. White scale bars: 1000 μm. Red scale bars: 200 μm. Numbers on both sides of the panels indicate dataset sizes (1^st^ row) and processing times (2^nd^-4^th^ rows) for voxel-based (left side) and APR-based (right side) versions of the workflow.

The PV+ channel was then segmented using a random forest classifier trained on sparsely annotated data directly on the particles, leading to a >60 times increase in the processing speed compared to a similar analysis performed on the voxel representation (Fig. 1h-j, Extended Data Fig. 1). Unlike existing histology pipelines^7,8^, segmentation is independently done on each acquired tile so no merging step is required, and the segmentation can be done during the acquisition of the next tile. The advantage of this approach is that it can scale up to an arbitrary large number of tiles and to parallel processing. Segmented objects on overlapping areas are therefore present on multiple tiles (2 or 4). By embedding each object in a feature space containing the object position and size, and using Lowe criteria^9^, duplicated objects are removed keeping the one with the higher SNR. We validated this approach on a synthetic dataset where the total number of objects was known: all objects were correctly matched even for large object densities (Extended Data Fig. 2).

The mouse brain sample was then registered to the Allen Mouse Brain Atlas at 25*μm* resolution using the voxel-reconstructed auto-fluorescence channel (Fig. 1k-m). Because only a lower resolution is needed for registration, APRs were lazily reconstructed at atlas resolution (thanks to their native pyramidal multi-resolution nature), and the registration step took only a few seconds as opposed to several hours previously required. After atlas registration, the density of PV+ neurons was measured in brain regions of interest. The cell distribution was found to be similar to previous studies^10^ (Fig. 1m).

To show the applicability of the proposed APR-native pipeline to samples relevant to human pathology, we imaged a human brain block of 1.7×2.7×0.3 cm from a patient with Alzheimer’s disease (AD) using an open-top light-sheet microscope^11^ (Supp. Table 1). The sample was stained for microglia and for amyloid *β* (A*β*) plaques (see methods), and it was imaged (17×12 tiles) in 3 channels, including an autofluorescence channel (Fig.2a-e). The A*β* plaques channel was used to perform the stitching and segmented using the same strategy as described above for the mouse brain. Autofluorescence remaining on the A*β* plaques channel after APR conversion was reduced before segmentation by subtracting the histogram-matched autofluorescence channel (see methods, Extended Data Fig. 3a-c). The stitched channel was used as a reference, and the two other channels were registered tile-by-tile to compensate for chromatic aberration, allowing the precise co-localization of different labels (Extended Data Fig. 3d-e). Using the segmentation results, it was then possible to compute the density (Fig. 2f-g) and the cortical depth of each segmented object by finding the closest sample margin (Extended Data Fig. 3f). Overall, we obtained a data-size reduction (117 times) and processing speedup (66 times) on the same order of magnitude as with the mouse sample. To demonstrate that pAPRica can scale to petabyte datasets a synthetic dataset of 1126 TB was created and processed and the same processing speed-up as the previous datasets was observed (see methods).

**Figure 2:**
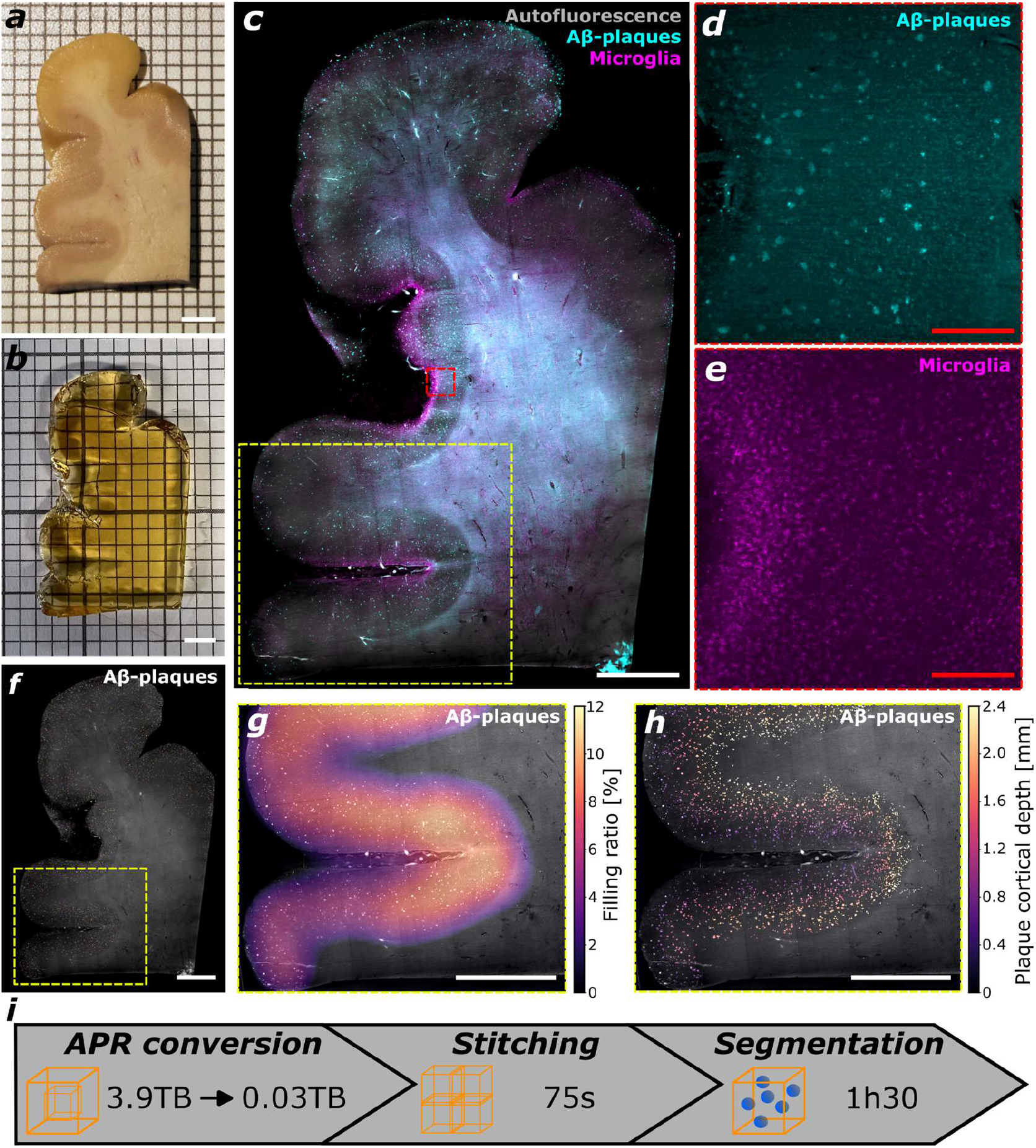
Processing of a large pathological human brain sample. ***a***, Image of a brain sample from a patient with Alzheimer’s disease before and ***b*,** after tissue clearing. ***c*,** Microscopic image of the whole sample after stitching of all channels (autofluorescence, Aβ plaques, microglia). The yellow dotted box delineates the subset displayed in (d),(e). ***d*,** Magnified view of A*β* plaques. **e,** Magnified view of microglia. ***f*,** Microscopic image with segmented (random colored) Aβ plaques. The yellow dotted box delineates the subset displayed in (d),(e). ***g*,** Aβ plaque density (volume filling ratio) highlighting their preferential location in the gray matter. ***h*,** Aβ plaques segmented and color-coded for their cortical depth. ***i*,** Pipeline running times and compression performance on the entire human brain sample. White scale bars: 4000 *μm*. Red scale bars: 400 *μm*.

Segmentation performance was benchmarked against the same pipeline run on raw voxel data and on voxel reconstruction of the APR (Extended Data Fig. 4). All three segmentations yielded comparable results (99.92% matching objects with an average center-of-mass deviation of 0.2 *μm*), suggesting that using the APR does not affect image-analysis accuracy while providing an important speed improvement. Further, we measured segmentation performance using the proposed random forest classification on APRs with different numbers of particles, obtained by varying the conversion parameters (Extended Data Fig. 5). The number of segmented objects and their positions remained stable when increasing the compression ratio. A drop in the Dice index was, however, observed when particles at native resolution were lost, corresponding to entirely downsampling the data. This suggests that information at full spatial resolution is crucial (Extended Data Fig. 5). APR conversion parameters are therefore automatically chosen such that this information is retained, except for empty tiles.

Altogether, we presented a scalable image-analysis pipeline that addresses the challenge of efficiently dealing with very large datasets, as they occur in next-generation histology. In addition to saving memory and storage, the presented APR-native pipeline considerably accelerates image processing. It thus enables whole-organ cell atlasing projects and the use of volumetric imaging in clinical pathology, opening the door to studying biomarkers in large 3D human samples.

## Methods

### PV+ mouse brain preparation and imaging

A mouse brain expressing the red fluorescent protein tdTomato in parvalbumin positive neurons (PV-TdTomato mice) was used for imaging. It was fixed with 4% PFA and cleared with the CLARITY protocol^22^, using the X-CLARITY*™* system. It was then immersed in a refractive index matching solution (RIMS) containing Histodenz (Sigma Aldrich) for at least 24 hours before being imaged. COLM imaging was performed by illuminating the sample by one of the two digitally scanned light sheets, using 488 *nm* and 561 *nm* laser lines. Emitted fluorescence was collected by a 4X objective (Olympus XLFLUOR4X N.A. 0.28), filtered at 525/50 nm and 609/54 nm (Semrock BrightLine HC) and imaged on an Orca-Flash 4.0 LT digital CMOS camera at 4 fps, in rolling shutter mode. A self-adaptive positioning system of the light sheets across z-stacks acquisition ensured optimal image quality over the whole sample thickness. Z-stacks were acquired at 3 *μm* spacing with a zoom of 4x resulting in an in-plane pixel size of 1.4 *μm* (2048×2048 pixels). Tiling was performed with a 40% overlap (409 pixels). The sample size was 16.54 mm in width, 21.63 mm in height and 6.44 mm in depth resulting in 2.4 TB of data.

### Human AD brain sample preparation and imaging

A fixed human brain sample was obtained from the Geneva Brain Collection^23^ in agreement with the Ethics Committee of the Canton of Geneva. The diagnosis of Alzheimer’s Disease (AD) was established on this sample by a neuropathologist.

The tissue was processed using a modified iDISCO protocol^24^. It was dehydrated with increasing concentrations of methanol (50%, 80%, 100%, 100%) at room temperature. It was then bleached with 5% *H_2_O_2_* overnight and rehydrated. Two washes of 1h in PBS/Triton X-100 (SIGMA T9284) were performed before permeabilization for 3 days at 37°C with 3% Triton, 0.3M glycine (Pan Reac applichem A1067), 5% DMSO (SIGMA 41640) and Saponin 10 *mg*/*mL* (SIGMA S4521). After permeabilization, the sample was blocked with gelatin (VWR-24350.262) for 2 days at 37°C and incubated with primary antibodies against Iba-1 (rabbit, 1:500, Wako #019-19741) and Aβ1-16 (mouse, 1:500, clone DE2, Merk Milipore MAB5206) for 7 days at 37°C in PBS with Tween-20 0,05% (Pan Reac applichem A4974), DMSO 5% and Heparin. After a series of washes in PBS/Tween the sample was incubated in secondary antibodies: donkey anti-mouse AlexaFluor 568 and donkey antirabbit AlexaFluor 647 (both 1:500, Jackson ImmunoResearch) at 37°C for 7 days in the same solution. Finally, it was washed in PBS/Tween for 1 day before clearing. Samples were cleared with gradual dehydration steps in methanol (50%, 80%, 100%, 100%), followed by a 3-hour incubation in 66% dichloromethane (DCM)/ 33% methanol at room temperature, and two 15-min washes in 100% DCM before immersion in Dibenzyl ether (DBE).

Imaging was done with an open-top light sheet microscope (ClearScope, MBF Bioscience) using double illumination provided by two digitally scanned light sheets, using 488 nm, 561 nm and 647 nm laser lines. Emitted fluorescence was collected by a 4X objective (Olympus XLFLUOR4X, N.A. 0.28), filtered (ET 405/488/561/640 nm Laser Quad Band Set, Chroma) and imaged on an Orca-Flash 4.0 LT digital CMOS camera at 4 fps, in rolling shutter mode. A linear adaptive calibration procedure objective was performed on the sample prior to imaging by aligning the center of the light sheets with the focal plane of the detection to correct for the refractive index inhomogeneity in the depth of the sample. Z-stacks were acquired at 4 *μm* spacing with a zoom of 4x, resulting in an inplane pixel size of 1.02 *μm* (2048×2048 pixels). Tiling was performed with a 10% overlap (102 pixels). The sample size after clearing and staining was 17.2 mm in width, 27.3 mm in height and 3.05 mm in depth resulting in 3.9 TB of data.

### Adaptive particle representation

The Adaptive Particle Representation (APR) is a content-adaptive image representation designed for large, sparse images and image stacks. It works by locally adapting the sampling density (voxel resolution) based on signal contents. This enables aggressive approximation of regions of low intensity variation, while maintaining the full voxel resolution in information-rich areas, such as edges. The adaptive sampling is determined in a globally optimal way with respect to a reconstruction condition, guaranteeing that the signal can be reconstructed with a bounded error at all original voxel locations. More precisely,

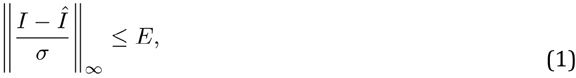

where *I* and *Î* are the original and reconstructed images, *E* is the user-defined error threshold, and *σ* (called the *local intensity scale*) is a space-dependent normalization function that guarantees equal adaptation to regions of different absolute brightness. It corresponds to a smoothed and rescaled estimate of the local standard deviation about each voxel.

APR particles form an octree structure in image space with square/cubic tree cells. Thus, if the voxel image is viewed as the leaf nodes of a full octree, an APR is composed of the leaf nodes of a pruned octree. Particles are placed at the center of each resulting cubic grid cell. The APR tree cells define a *resolution function*, *R*(**x**), which at each point **x** in the image takes the value of the side length of the APR cell containing that point. Concretely, since the APR uses dyadic refinement, if the point **x** is contained in a grid cell at level 0 ≤ *l* ≤ *l*_max_ in the tree, the resolution function takes the value *R*(**x**) = 2^*l*_max_-*l*^. Here, level 0 is the root of the tree, and cells at level l_max_ correspond to the original voxels. A defining feature of the APR is that it allows the reconstructed value (*Î* in Eq. 1) at any location **x** to be any positively weighted average of particles within a radius *R*(**x**) of **x**. For any such reconstruction formula, Eq. 1 holds. However, in order to determine the globally optimal APR of an image in linear time, Eq. 1 is not used directly. Instead, the problem is reformulated as a bound on the resolution function

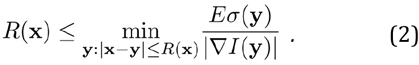

Thus, computing the APR of an image requires determining the local intensity scale *σ* as well as the gradient magnitude |∇*I*| at all original voxel locations. Exactly how these quantities are computed is a design choice. Here, we used the APR library *LibAPR*^12^, where both the gradient field and the local intensity scale are computed from a smoothing cubic B-spline representation of the original image. Based on Eq. 2, the APR is then computed in a globally optimal way, i.e., no other multi-resolution representation of the same image can exist that would fulfill Eq. 1 with less sampling points. This optimality and guaranteed approximation bound are the defining features that distinguish the APR from super-voxel approaches.

The computational savings afforded by an APR representation of an image scale with the ratio between the number of voxels in the original image and the number of particles in the APR. This ratio is called the *Computational Ratio* (CR) and can be understood as the processing speedup for linear time-complexity algorithms. The ratio between the size of the uncompressed input image file in Bytes and the size of the compressed APR file in Bytes is called the *Memory Compression Ratio* (MCR). It is always higher than the CR because particles are losslessly compressed using C-Blosc when saved on non-volatile memory.

We refer the reader to Cheeseman et al.^5^ and its Supplement for implementation details and mathematical proofs of the theory behind the APR, and to Jonsson et al.^6^ for additional information on efficient APR data structures and APR-native processing algorithms such as convolutions and filters.

### Automatic APR conversion

Converting a 3D image stack to an APR requires several parameters to be set. Many of these are specific to the microscope setup or can reliably be kept constant within a given experiment. This includes the voxel size in each dimension, the error bound *E* in Eq. 1, as well as the degree of smoothing in the B-spline approximation. However, three threshold parameters (see Supp. Table 2) are introduced to allow more direct control of the APR adaptation. These depend on the signal content, such that optimal values vary between different regions of a sample.

Zero gradients are taken to mean that the resolution function is unrestricted. Thus, *I_th_* and ∇*_th_* are hard thresholds with the effect that gradient-computation errors are ignored in regions of low intensity or low gradient magnitude. The intensity scale threshold *σ_th_* sets a minimum value for the intensity scale across the image. This is useful to avoid adaptation to noise due to normalization of gradients in flat regions, where the intensity scale tends to the average noise range. Together, these three thresholds serve the same purpose of separating useful signals from uninformative regions, such that the latter can be aggressively compressed. However, care must be taken as excessively large values of any one threshold can result in loss of image content.

While optimal threshold values are image-dependent and somewhat subjective to the task, manual tuning is not feasible in experiments of the scale considered here. Thus, we have devised a simple yet robust scheme to automatically find suitable parameter values. The intensity threshold *I_th_* is set manually to a conservative value based on the background intensity of the optical setup, to filter out regions outside the sample (but not necessarily background within the sample). The remaining thresholds are then computed automatically by first uniformly subsampling the computed gradient and local intensity scale values in regions where *I* > *I_th_* and then applying the iterative minimum crossentropy thresholding algorithm by Li et al.^13^ to histograms of the two fields to determine the thresholds.

When used at acquisition time, the pipeline can automatically run quality checks after conversion and discard the raw voxel data to save space.

### Volume stitching

Samples are imaged by the microscope tile by tile using a raster scan pattern where each tile is a 3D array sequentially acquired. Our pipeline supports sparse tiling (i.e. some tiles can be missing if, e.g., a pre-scan was performed to not scan empty tiles), as long as each tile is connected to the others by at least one of its sides. This was adapted for APR use by drawing inspiration from previous work^14^.

First, the neighborhood map is computed for each tile, consisting of the edge-connected neighbor tiles of that tile. Then, for each pair of neighboring tiles the displacements for aligning those tiles (*d_x_*, *d_y_*, *d_z_*) are computed using the phase cross-correlation^15^ of the maximum-intensity projections (*I_x_*, *I_y_*, *I_z_*). If artifacts are present in the volume (e.g. air bubbles embedded around the sample), a masked version of the phase cross-correlation can be used^16^. The correlation function can be changed by users if needed. When used for batch processing, as opposed to online processing during acquisition, the maximum-intensity projections are pre-computed (only on the expected overlap areas to remove noise in the phase cross-correlation) to enable the stitching to be performed with a single reading of each tile. When used for online processing, the maximum-intensity projection is computed as soon as a tile has been completely acquired and saved for when new neighboring tiles are acquired. If desired, maximum-intensity projections can be limited to a certain range of depths. In this paper, we computed all maximum-intensity projections over entire stacks.

Doing that, 6 displacements (2 for *x*, 2 for *y* and 2 for *z*) are determined for each pair of neighboring tiles and the most reliable one is kept. For dense labeling, the reliability is computed as the leastsquares difference between registered maximum-intensity projections. For sparse labeling, the reliability is computed as the ratio between the maximum of the sum of the registered areas and the sum of both individual maxima (if the registration is correct, then maximas are aligned and the ratio should be close to one). The reliability function was always penalized by multiplying the reliability by the square root of the mean signal in the registered areas, so that empty tiles (or tiles with fewer signals) are given a lower reliability and are discarded during the global optimization step.

The displacements are stored in three graphs (one for each dimension), where each vertex corresponds to a tile position and each edge corresponds to a displacement between neighboring tiles. A corresponding reliability graph is also constructed for each dimension. Finally, each displacement graph is globally optimized using the maximum spanning tree of the corresponding reliability graph to satisfy the constraint that each loop in the graph should sum to 0.

All of these steps are directly computed on the APR with the exception of the phase cross-correlation. For this step, the maximum-intensity projection is computed on APR, from which a 2D pixel image is then reconstructed to compute the phase cross-correlation. Reverting to pixels for this step is not limiting, because it is only in 2D on a per-tile basis and therefore has a low memory footprint. Finally, stitched tile positions are stored in a database so they can later be used to compute segmented object positions, to merge data, and for visualization.

Stitching speed was compared with TeraStitcher^14^ without the final step of merging the data. Multiple instances of TeraStitcher were run at the same time using OpenMP. The best performance was obtained when using 4 CPU cores. We created a synthetic data set consisting of 4×4 tiles of 512×512×2048 uint16 voxels (around 2.2 GB/tile) with various computational ratios simulated by varying the number of objects in the images (3D spheres of random diameters with poisson noise). Voxel data was converted to APR using automatic parameter determination (see above).

For the human AD brain sample, channels were co-registered to compensate for light-sheet misalignment, especially along the depth axis (Extended Data Fig. 3d,e). To do so, the Aβ-plaque channel was used as a reference, and the two other channels were registered tile by tile using a dedicated module of the proposed pipeline.

### Object segmentation

For segmenting structures of interest, such as labeled cells, a particle classification strategy was used. 3D filters (gradients, laplacian, gaussian, difference of gaussian and particle level) are applied directly on the APR data. A random forest with 100 estimators^17^ was trained on sparsely annotated data taking as input the intensity data and the filtered data.

The proposed pipeline does not merge tiles before performing. Therefore, segmentation can be done immediately after a tile has been completely acquired, while the next is being acquired, and in parallel for different tiles.

Features (such as diameter, volume, eccentricity, etc.) and positions of segmented objects were extracted, and a merging strategy inspired by Lowe^9^ was implemented to merge the high-level information without actually merging the APR data. For each pair of neighboring tiles, the two nearest neighbors (NN) of each object in the overlap area of the first tile are found from amongst the objects in the overlap area of the second tile. If the ratio between the first NN and the second NN is smaller than 0.7, and if the object centers are closer than a threshold distance (typically a quarter of the object size; the reliability estimation of the tile position can also be used to adapt this threshold), then objects are considered to be the same and merged. Objects that are touching an edge of an overlap area are discarded to avoid counting them twice. This strategy was benchmarked on a synthetic data set (Extended Data Fig. 2), where the correct number of objects was extracted under random displacements between tiles.

The proposed merging strategy was developed for objects that don’t span on multiple tiles. It allows to estimate cell densities and

If the random forest can not be trained prior to acquiring the data, the filter can still be computed and saved during acquisition. Because filter computation is the bottleneck for image segmentation (it consists of applying 3D convolution filters on the APR^6^), this is going to accelerate later offline training and use of the segmentation model.

The outputs of the classification step (probability map, segmentation mask, connected components) were saved in the same file as the original APR, avoiding the need to save the tree data structure twice. If desired, however, they can also be saved in different files.

For the sake of comparison, segmentation of one tile from the PV brain dataset was also performed on the raw voxel data and on voxel data reconstructed from the APR. The voxel segmentations were performed using the ZEISS Arivis Software. Background subtraction (computed as the Gaussian blurred image with sigma = 213 voxels) was followed by denoising using a Gaussian filter of size 25 *μm* and finally by blob detection (22 *μm* maximum diameter, probability threshold 12.5%, split sensitivity 75.9%). Segmentation results were then compared by matching segmented objects using the same approach as described above: 99.92% of segmented cells were matching with an average center-of-mass mismatch of 0.2 *μm* and an equivalent average segmented object diameter of 13 *μm* (Extended Data Fig. 4). This experiment suggests that converting images to APR is not detrimental to segmentation performance.

The influence of the CR (defined above under “Adaptive particle representation”) on segmentation results was studied by generating different APRs with increasing error threshold *E*. Increasing the error threshold allows the resolution function to assume larger values uniformly for each voxel, see Eq. 2, hence increasing the CR of the resulting APR. For each obtained APR, a random forest segmentation model was independently trained on the same manual annotations to segment plaques in the human brain dataset. The number of segmented objects and the Dice index (with the lowest CR as reference) were computed for each APR (Extended Data Fig. 5). See main text for interpretation.

### Volume merging

Certain applications, such as registering a sample to an atlas, require a merged volume. Fortunately, these applications usually require a lower sample resolution, avoiding the need to reconstruct the original data footprint. The pyramidal multi-resolution structure of the APR enables efficient reconstruction at power-of-two reductions of the original voxel resolution. Moreover, by storing coarsened particle values (referred to as *APR Tree* in 6), the reconstruction can be done in a lazy manner, avoiding the need for reading the high-resolution information. This allowed merging the present APR brain data within seconds, whereas merging 72 tiles on raw voxels took 2h 10s.

Reconstructions at finer resolutions can be obtained by interpolation from the next higher power-of-two APR level. Tiles are then merged using any merging strategy, such as taking the average or the maximum (used for the figures in this paper) of the overlapping areas. More complex blending strategies can also be used. Preprocessing (e.g. histogram equalization) can be performed directly on the APR, prior to voxel reconstruction and merging, in order to benefit from the computation speedup.

### Registration to an Atlas

The merged mouse brain volume was registered to the 25 *μm* resolution Allen Mouse Brain Atlas^18^ using the AMAP pipeline^19^ through the Brainreg API^20^.

### Visualization

A Napari^21^ plugin was developed for displaying and converting APR volume images. It allows for opening and displaying APR files with a simple drag-and-drop user interface. Napari requires an object with a slicing property and thus allows reconstructing only the current frame in pixels. This approach enabled benefiting from Napari’s displaying capabilities (multi-tile, 3D blending, etc.) out of the box.

A lazy APR loader is also provided, which allows reading a sub-volume of an APR without a need for loading its full volume into memory. This permits reconstructing sections of large samples with hundreds of tiles in just a few seconds.

It is also possible to reconstruct a 3D volume at higher resolution in just a few seconds, thanks to the tree structure of the APR and similarly to pyramidal files.

### Segmented A*β* plaque density and cortical depth estimation

Density (volume filling ratio) of Aβ plaques was estimated using a scaled Gaussian kernel of size 200 *μm* on the segmentation mask. Aβ plaque depths were computed for their intensity-weighted centroid. First, sample borders were extracted using the averaged auto-fluorescence channels after applying a gradient filter in both *x* and *y* directions followed by a threshold (determined manually). The top and bottom edges were removed to avoid biasing the analysis. The obtained sample border mask was then sub-sampled by a factor 8, and voxels belonging to the sample borders were gathered in a (*N x* 3) matrix. For each segmented Aβ plaque, its depth was computed as the Euclidean distance to the nearest neighbor between the plaque centroid coordinates and the sample margin coordinate matrix. Figure 2h with color-coded segmented objects was obtained by linearly interpolating the hue from purple (shallow depth) to yellow (large depth) from color look-up tables.

### Petascale synthetic dataset creation and processing

A 256×256 tiles dataset was created with each tile being a (2048, 2048, 2048) pixel array. Each tile was randomly populated with objects of random radius uniformly distributed from 0 to 20 pixels with a density of 20 000 objects per tile in order to obtain a similar CR as the previous acquisitions. Naively generating and converting to APR such a large dataset would take approximately 38 days (80% dedicated for generating the voxel data and 20% dedicated for converting to APR and saving) which is a waste of resources for a synthetic experiment. The dataset was therefore generated by creating a single (2048, 2048, 2048) tile with 20 000 objects and with periodic conditions at the 4 connected edges regarding object positions which mimics a 20% overlap in a tiled acquisitions. This tile was then converted to APR (Poisson noise was directly added on APR) and copied and pasted 16536 times to obtain a 256×256 dataset with CR of 73 and MCR of 139. As opposed to other samples, the APR data was saved on a 12 TB HDD because it would not fit on our workstation SSD so the reading and writing speeds were slower. This dataset was stitched in 24h30min. Reconstructing the whole dataset downsampled by a factor 32 took 6h30min and we estimated the segmentation time to be 577h, or 24 days, by segmenting 3×3 tiles and extrapolating to the whole dataset. Acquiring such a large dataset would take more than a year with the microscope working 24h/24 everyday.

### Computing hardware

The workstation used for analysis was custom made. The motherboard is a Z590 extrem model hosting a i9-11900KF CPU with 128 GB of 3200 MHz DDR4 RAM. Raw data were saved on 12 TB WD Gold HDD and APR were saved on 4TB Samsung 870 EVO SSD (except for the petascale synthetic dataset). The total cost of the workstation in 2022 is less than 3000$.

## Data availability

The study data are available from the corresponding author upon request.

## Code availability

Code will be made available on GitHub upon submission. The pipeline API is written in Python and we also provide a Napari plugin to use most of the pipeline functionalities with a graphical user interface.

## Acknowledgements

We thank Dr Enikö Kövari for kindly selecting and providing human brain tissue from the Geneva Brain Bank. This work was supported in parts by the Center for Scalable Data Analytics and Artificial Intelligence (ScaDS.AI) Dresden/Leipzig, funded by the German Federal Ministry of Education and Research (BMBF, Bundesministerium für Bildung und Forschung).

## Author contributions

J.S., J.J., C.M.L. and I.F.S. designed the research. C.M.L. and I.F.S. conceived the project. J.S., J.J., T.J.S., I.G., L.B., B.L.C. and S.P. performed research. J.S. built the workstation and analyzed data. J.S., J.J., C.M.L. and I.F.S. wrote the paper, with input from all authors. All authors approved the manuscript.

## Competing interests

J.S., J.J., B.L.C. and I.F.S. have a provisional patent application related to the proposed pipeline.

