## Supplementary Material for "Efficient image analysis for large-scale next generation histopathology using pAPRica"

**This file includes:**

Supp. Tables 1-2
Extended Data Figures 1-5

| Datasets | Voxel size (x,y,z) | *nchannel* | *ntiles* | Size | CR* | MCR** | Stitching | Segmentation |
|---|---|---|---|---|---|---|---|---|
| PV mouse brain | (1.4, 1.4, 3) $\mu m$ | 2 | 72 | 2.4 TB | 79 | 115 | 55 $s$ | 1$h$ 18$min$*24 |
| Human AD brain sample | (1, 1, 4) $\mu m$ | 3 | 204 | 3.9 TB | 66 | 117 | 75 $s$ | 1$h$ 30$min$ |
| Synthetic petascale | NA | 1 | 65536 | 1126 TB | 73 | 139 | 24h30 | 577h |

**Supp. Table 1: Dataset specifications and APR processing parameters.** *The Computational Ratio (CR) is the ratio between the total number of voxels and the total number of particles for the entire dataset. **The Memory Compression Ratio (MCR) is the ratio between the size of the raw image file (uncompressed tiff file) and the size of the APR file (losslessly compressed using C-Blosc) in Bytes.

| Symbol | Name | Description |
|---|---|---|
| $I_{th}$ | Intensity threshold | Gradients are considered 0 where $I < I_{th}$ |
| $\nabla_{th}$ | Gradient threshold | Gradients with magnitude smaller than $\nabla_{th}$ are considered 0 |
| $\sigma_{th}$ | Intensity scale threshold | Local intensity scale is truncated from below to $\sigma_{th}$ |

**Supp. Table 2: Threshold parameters used to control the APR conversion**.

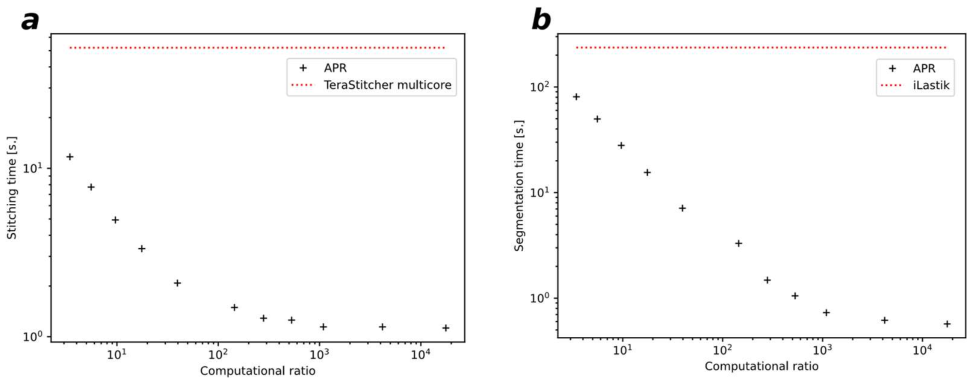


**Extended Data Figure 1: Benchmarks.** ***a,*** APR stitching times (black crosses) for different Computational Ratios (CR), compared with TeraStitcher on raw voxels (dotted red line). 4×4 tiles each of size 512x512x2048 were stitched using both methods. **b,** APR segmentation times for different CR (crosses), compared to the iLastik segmentation time (dotted line) for one 512x512x2048 tile on raw voxels.

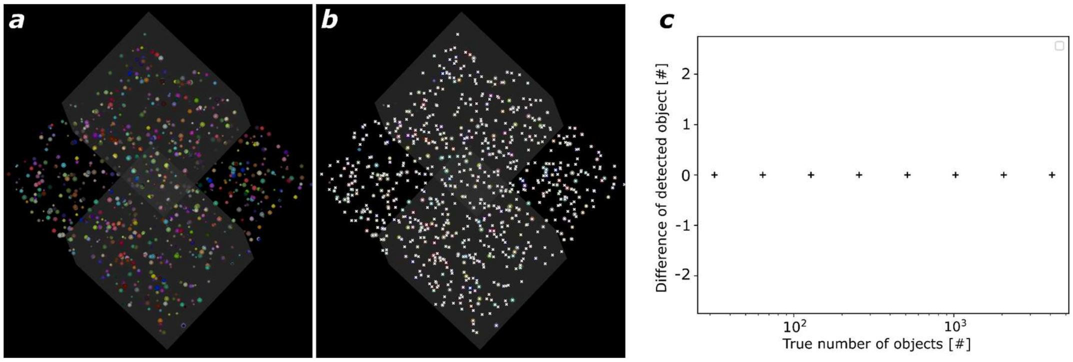


**Extended Data Figure 2: Merging segmented objects on multi-tile data.** ***a***, Representation of a synthetic 2x2 tiles data set, created with various numbers of randomly placed objects. A random displacement was added to each tile as would happen during a real microscopy acquisition. Each tile was individually segmented, leading to multiple occurrences of the same object in overlap areas**. *b,*** Representation of the merged segmentation with *o*bjects consolidated in overlapping areas after computing the correct registration (see methods section "Object segmentation"). ***c,*** Plot of the differences between the numbers of detected objects after merging and the true number of objects in the synthetic data set, as a function of the true number of objects. Each measurement was repeated on 128 independent randomly generated data sets; the observed difference was always 0.

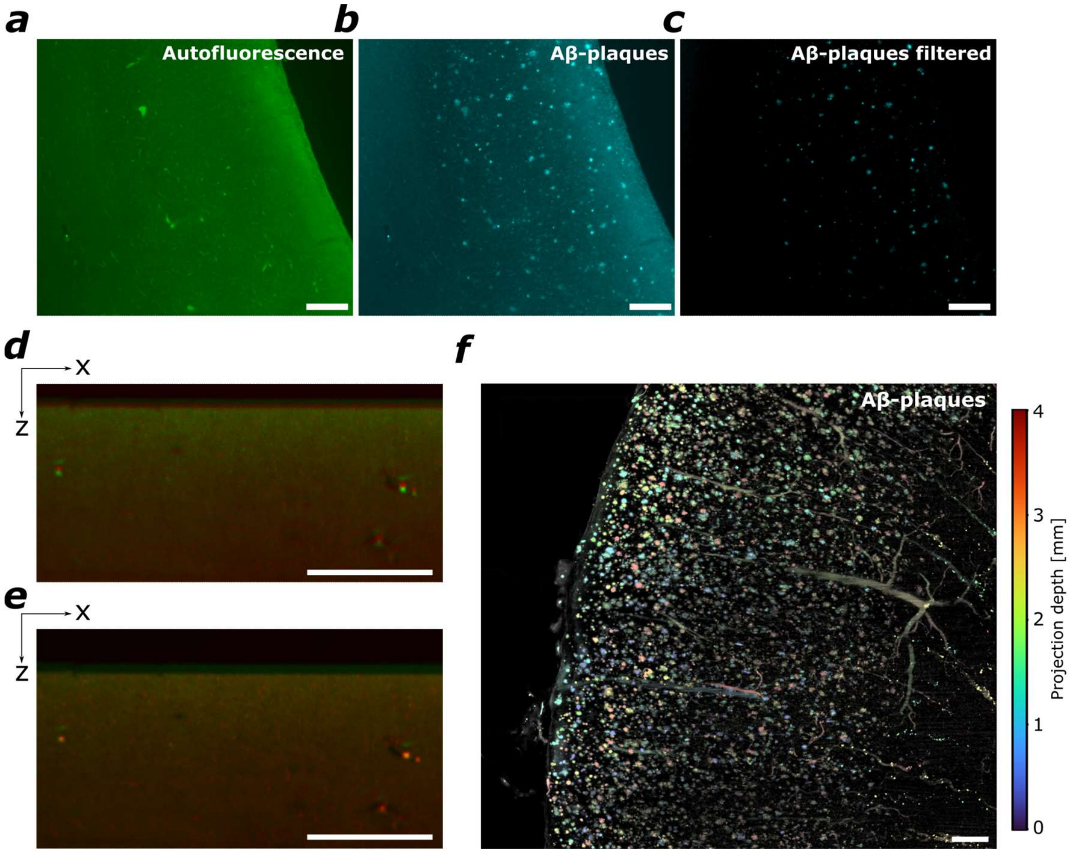


**Extended Data Figure 3: Segmentation of Aβ plaques in a human brain sample with Alzheimer's Disease.** ***a,*** Autofluorescence channel. ***b,*** Aβ plaques. **c,** Filtered image form (b): to remove spurious detections due to autofluorescence, the autofluorescence channel was subtracted from the Aβ-plaque channel after histogram matching. ***d,*** Merged autofluorescence and Aβ-plaque channels before intra-channel registration. **e,** Same image as in (d) after intra-channel registration to correct for chromatic aberration. ***f,*** Segmented Aβ plaques color-coded for projection depth, highlighting the 3D distribution of plaques. Segmentation was performed directly on the APR.

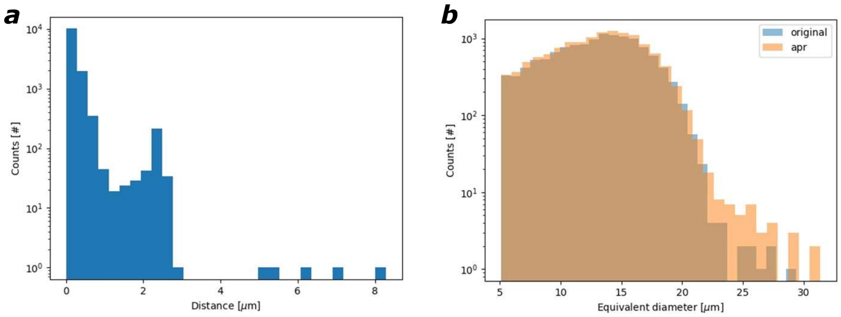


**Extended Data Figure 4: Comparison of segmentations performed on raw voxels and on reconstructed APR data.** ***a,*** Segmentations were performed using the same pipeline in the Arivis software on the raw-voxel PV brain dataset and on the corresponding voxel data reconstructed from the APR. The bar plot shows the histogram of the center-of-mass distances between segmented objects after matching them; the average distance is $0.2\ \mu m$. ***b,*** Histogram of equivalent diameters (volume cube root) for objects detected in the raw data and in the reconstructed data after conversion to APR. 1.6% more objects were segmented on the later, because the APR also acts as a denoising algorithm, leading to improved signal-to-noise ratio (SNR) and better segmentation of dimmer objects even after converting back from APR to voxels.

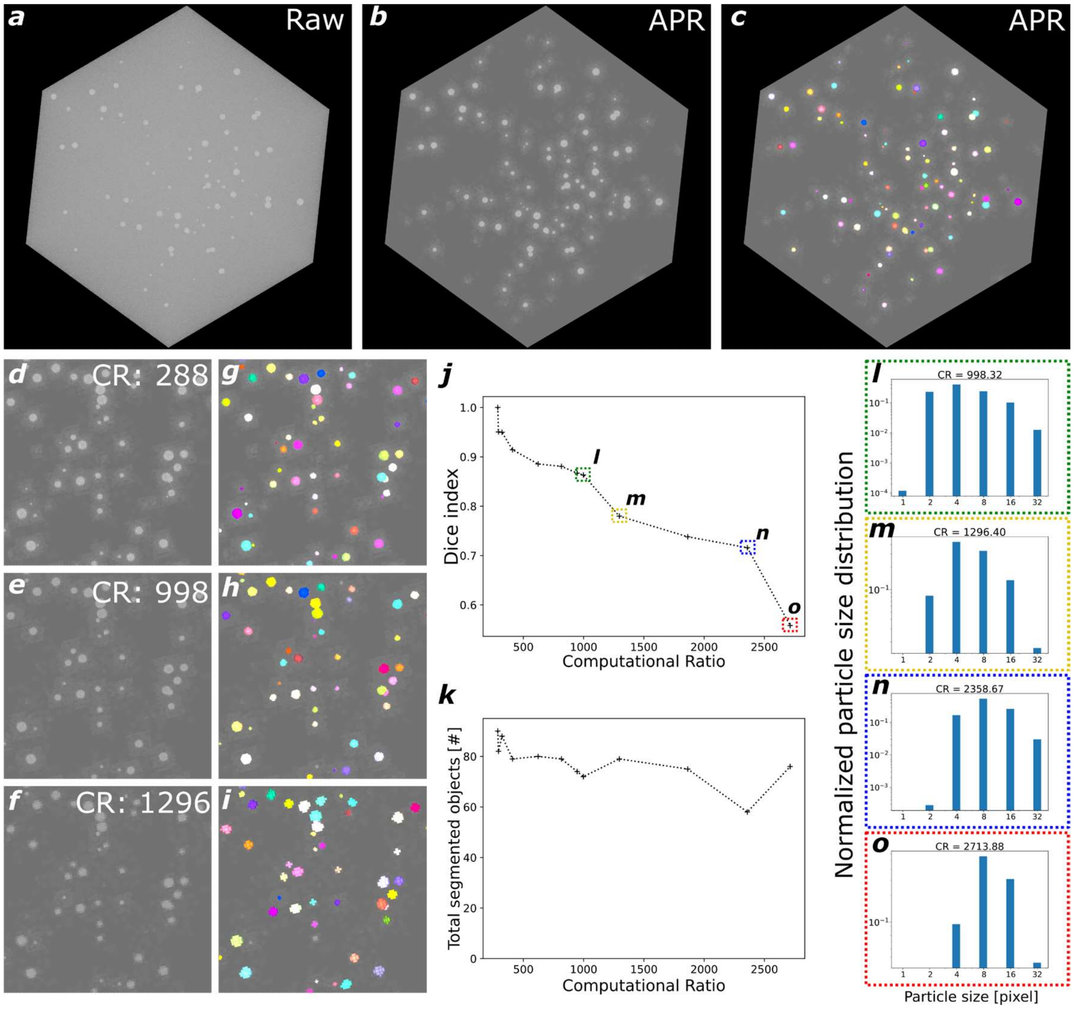


**Extended Data Figure 5: Segmentation with increasing CR.** ***a,*** Synthetic dataset consisting of spheres of random sizes and locations with SNR=10. ***b,*** Same dataset as in (a) after conversion to APR. The lookup table was kept the same as in (a), which highlights the denoising and background removal capability of APR. ***c,*** Same dataset as in (b) with the segmentation result overlaid in color. ***d, e, f,*** Magnified 3D rendering of the same dataset for different APR error thresholds $E$ (Eq. 1), leading to different CR values. ***g, h, i,*** results of segmentations performed on (e), (f) and (g) displayed as color overlays. Annotations used to train the random forest were kept the same, but the random forest was retrained on each APR of different CR. ***j, k,*** Dice index and number of segmented objects, respectively, for different CR (computational ratio) values. All segmentations were performed on the 3D APRs. ***l, m, n, o,*** APR particle size histograms corresponding to the colored boxes in (j). Each strong decrease in Dice index in (j, between the green and yellow, as well as between the blue and red box) corresponds to losing particles at a given resolution, leading to downsampled data. Between each loss of a resolution level, a decreasing exponential behavior is observed before reaching a plateau. This can be interpreted as the APR retaining the most important information, allowing it to maintain a constant Dice index up to the point where all particles of highest resolution are lost, which causes a drop in the Dice index. Interestingly, after each drop, the number of detected objects increases. This could be because, when a region is downsampled, the particle values are computed as a mean, leading to an increase of the local SNR. This increase in SNR helps the random forest segment dimmer objects.
